# Dynamics and predicted drug response of a gene network linking dedifferentiation with β-catenin dysfunction in hepatocellular carcinoma

**DOI:** 10.1101/347666

**Authors:** Claude Gérard, Mickaël Di-Luoffo, Léolo Gonay, Stefano Caruso, Gabrielle Couchy, Axelle Loriot, Junyan Tao, Katarzyna Konobrocka, Sabine Cordi, Satdarshan P. Monga, Emmanuel Hanert, Jessica Zucman-Rossi, Frédéric P. Lemaigre

## Abstract

Alterations of individual genes variably affect development of hepatocellular carcinoma (HCC), prompting the need to characterize the function of tumor-promoting genes in the context of gene regulatory networks (GRN). Here, we identify a GRN which functionally links LIN28B-dependent dedifferentiation with dysfunction of *CTNNB1* (*β-CATENIN*). LIN28B and CTNNB1 form a functional GRN with SMARCA4 (BRG1), Let-7b, SOX9, TP53 and MYC. GRN activity is detected in HCC and gastrointestinal cancers; it negatively correlates with HCC prognosis and contributes to a transcriptomic profile typical of the proliferative class of HCC. Using data from The Cancer Genome Atlas and from transcriptomic, transfection and mouse transgenic experiments, we generated and validated a quantitative mathematical model of the GRN. The model predicts how the expression of GRN components changes when the expression of another GRN member varies or is inhibited by a pharmacological drug. The dynamics of GRN component expression reveal distinct cell states that can switch reversibly in normal condition, and irreversibly in HCC. We conclude that identification and modelling of the GRN provides insight into prognosis, mechanisms of tumor-promoting genes and response to pharmacological agents in HCC.

---

Hepatocellular carcinoma (HCC) is the most prevalent liver primary tumor and the third-most common cause of cancer death worldwide ^1, 2^. Liver resection, transplantation or radiofrequency ablation are curative therapeutic options, useful in only less than 30% of the cases. In advanced HCC, administration of Sorafenib or other drugs provides survival benefit to unresectable HCC, yet without curing the disease ^3^. Therefore, identification of new molecular strategies is urgently needed.

Various etiologies are associated with HCC, leading to significant heterogeneity in clinical outcome, histology, transcriptomic profile and mutational spectrum ^4, 5, 6, 7, 8^. Such heterogeneity can cause a variable response to therapeutic agents, as in mouse models with *Ctnnb1*-induced HCCs which show heterogeneous sensitivity to CTNNB1 (β-CATENIN) inhibitors ^9^. Thus, designing novel therapeutic strategies against HCC requires not only the identification of inhibitors of individual tumor-promoting genes but also the characterization of the molecular networks in which those genes exert their functions.

Dedifferentiation of hepatic cells contributes to HCC progression ^10, 11, 12^. In this context, poorly differentiated HCC develop as a result of forced induction of LIN28B, a RNA-binding protein which is repressed during normal hepatic cell differentiation. LIN28B is re-expressed in a subset of human HCC’s characterized by high serum levels of α-foetoprotein ^13, 14^, thereby associating dedifferentiation, HCC progression and LIN28B expression ^15^. In parallel, *CTNNB1* is one of the most frequently mutated genes and drivers of HCC, with a mutation rate of 11%-37% ^1^. Therefore, we here explore the possibility that HCC progression depends on a gene regulatory network (GRN) linking LIN28B-dependent dedifferentiation with CTNNB1 dysfunction.

Several approaches can be implemented to identify GRNs ^16^. We here select a method which captures the dynamics and biological logic of the system, and identify a HCC-promoting GRN comprising several members connecting LIN28B with CTNNB1 via Let-7b, MYC, SMARCA4 (BRG1), TP53 and SOX9. We further investigate the system-level dynamics of the GRN with help of a quantitative mathematical model, which is calibrated and validated using quantitative mRNA and protein expression data from HCC cell lines, patient databanks and mouse models. *In silico* simulation of the quantitative impact of a tumor-promoting gene on the expression of the other GRN members revealed the existence of distinct HCC cell states which may be at the origin of inter-tumoral heterogeneity.

Several components of the GRN identified here in HCC are instrumental in development of other cancers, and we provide evidence that the GRN is functional in several gastrointestinal cancers. Finally, we built a web-based platform which determines if the GRN is functional in individual HCC samples and allows *in silico* evaluation of potential anticancer strategies targeting the GRN.

## Results

### Identification of a gene regulatory network driving hepatocellular carcinoma

We selected an approach in which GRN members must meet two criteria. First, their role in tumor-promotion must be validated by animal experimentation and/or high-throughput sequencing data from patients. Second, GRN members fitting the first criterion must be connected by direct or indirect functional links characterized by protein-protein and protein-DNA interactions, or epistatic relationship identified in loss- and gain- of function analyses.

By combining data from the literature, we first reconstituted a GRN comprising 7 cross-regulating components: the miRNA *Let-7b,* the RNA-binding protein LIN28B, the ATP-dependent helicase SMARCA4, and the transcription factors SOX9, MYC, CTNNB1 and TP53 (Fig. 1a, left): Let-7 is a tumor suppressor whose maturation is repressed by re-expression of LIN28B in cancer, including in HCC ^17, 18, 19, 20^. LIN28B can cooperate with CTNNB1 to promote tumor development in mouse models of colorectal cancer ^21^, it is stimulated by MYC and its overexpression is sufficient to induce HCC in mice ^15^. In addition, MYC expression is stimulated by a CTNNB1/SMARCA4 complex, in which SMARCA4 is a chromatin regulator mutated in HCC ^6, 7, 22^. Lastly, TP53 and MYC regulate each other ^23^, and SOX9 is both a stimulator of CTNNB1 expression and target of Let-7b ^24, 25^. Thus, this literature-based GRN connects LIN28/Let-7, a key axis for cell differentiation, with CTNNB1 whose deregulation is often involved in HCC development. Within the *Let-7* family, *Let-7b* is here selected since it is the most differentially expressed family member between normal and HCC samples ^26^. The functional links in the GRN reflect direct or indirect regulations which were determined by transfection, gene inactivation, protein-protein and protein-DNA interaction studies ^8, 10, 11, 12^. When interactions have been identified in non-hepatic cells, we verified whether they also occurred in cultured HCC cell lines (see below).

**Figure 1.**
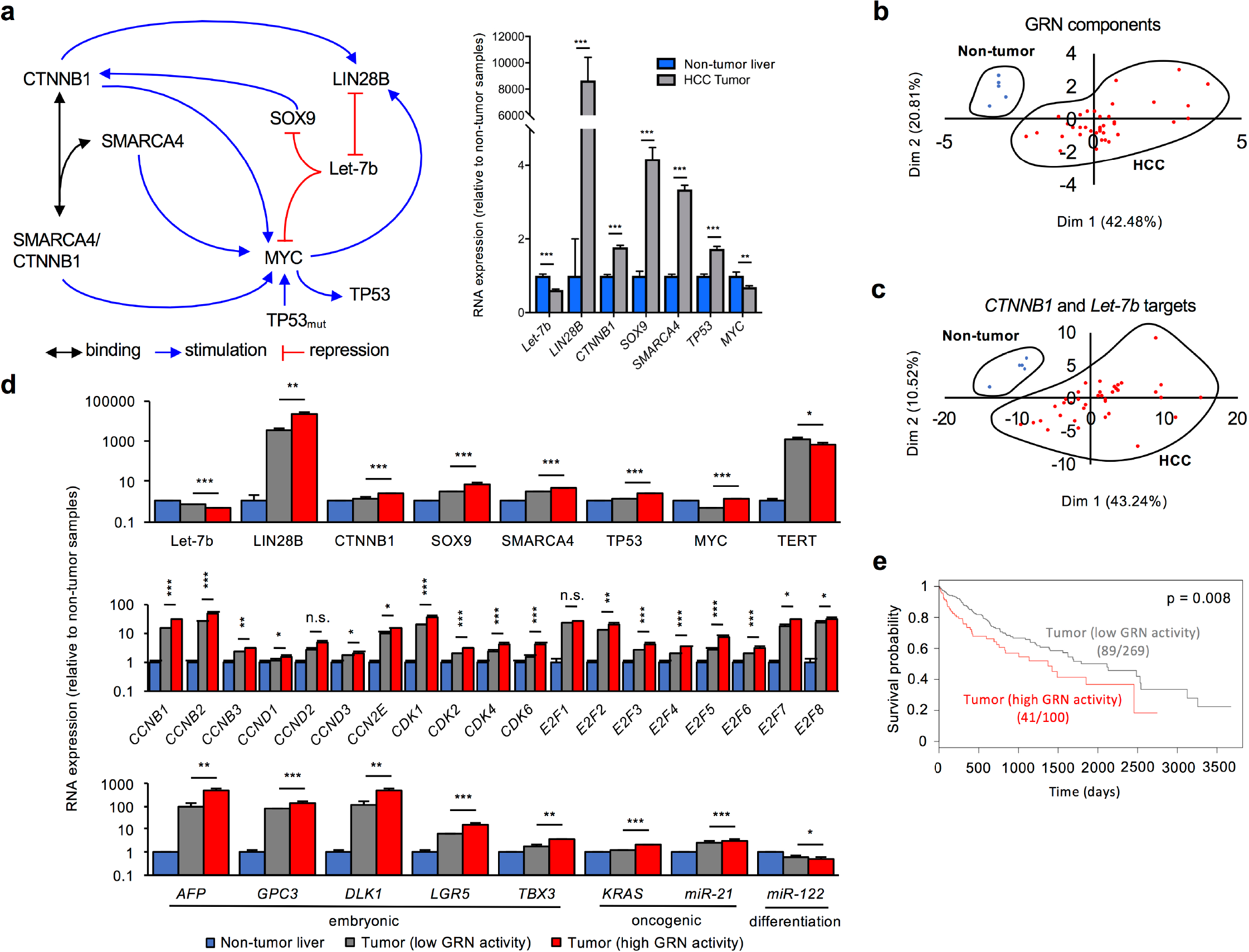
Identification of a GRN involved in HCC. (**a**) Structure of the GRN (left), and RNA levels of GRN components in normal (non tumor) tissue (n=50) and HCC tumors (n=369) from the HCC cohort in TCGA (right). (**b**) PCA plots based on the expression of the GRN components, and (**c**) on selected CTNNB1 and *Let-7b* targets (listed in Supplementary Table 1) in normal and HCC samples show that the activity of the GRN is increased in HCC. Each dot represents the mean of 10 samples grouped in alphabetic order of the sample ID. (**d**) Tumor samples were ranked according to low or high GRN activity (see text), and expression of the GRN components differed significantly between low and high GRN activity samples. Highest GRN activity (top) correlated with highest expression of proliferation (middle), embryonic and oncogenic markers (bottom), and lowest levels of the differentiation marker miR-122 (bottom). (**e**) GRN activity negatively correlated with vital prognosis of patients. Kaplan-Meier curves showed that patients with high GRN activity (median time = 1372 days, 41 dead patients amongst 100) exhibited a lower survival probability than patients with low GRN activity (median time = 2116 days, 89 dead patients amongst 269) (Wilcoxson test, p = 0.008).

Data from the The Cancer Genome Atlas (TCGA; Fig. 1a, right) show that expression of *Let-7b* is reduced in HCC as compared to adjacent non tumor liver tissue. This is consistent with the concomitant overexpression of LIN28B, a repressor of *Let-7b.* All other GRN components, except MYC, were increased in HCC. Further, a principal component analysis (PCA), which considers only the expression of the 7 GRN components in 50 non tumor controls and 369 HCCs revealed that non tumor tissue and HCC tumors clustered separately (Fig. 1b and Supplementary Fig. 1a).

We then considered the 100 HCC cases of the TCGA cohort with highest expression of a GRN component, and the 100 cases with lowest expression of the same component. We then verified whether the other GRN components were correlatively up- or down-regulated. Except for Let-7b, high expression of a component correlated with high expression of the other GRN components (Supplementary Fig. 2a). Also, in a PCA analysis the tumor samples with high expression of a GRN component clustered together and separate from the tumors with low expression and from the non tumor samples (Supplementary Fig. 2b). Together, these analyses suggest that the expression of the GRN components is correlated.

Importantly, PCA investigating multiple targets of *CTNNB1, Let-7b-3p* and *Let-7b-5p* (Supplementary Table 1) also identified separate clusters for non tumor tissue and HCC tumors, indicating that the components of the GRN were not only misexpressed in HCC but also that they were actively regulating their targets (Fig. 1c and Supplementary Fig. 1b). Therefore, we considered that concomitant overexpression of the GRN components LIN28B, SMARCA4, SOX9, CTNNB1 and TP53, and downregulation of Let-7b was indicative of GRN activity.

Out of the 369 HCCs from the TCGA cohort whose GRN component expression had been analyzed by PCA we selected the 150 samples with the highest Dimension 1 to which *CTNNB1, SMARCA4, SOX9, MYC* and *TP53* expression contribute the most, and out of the latter we selected the 100 samples with highest Dimension 2 to which *LIN28B* and *Let-7b* contribute the most (Supplementary Fig. 1c). These 100 samples were defined as displaying high GRN activity and were compared with the other 269 tumors, which were defined of low GRN activity, and with non tumor liver tissue (Fig. 1d). The expression of the GRN components differed significantly between the tumors with low and high GRN activity. Moreover, the samples with high GRN activity displayed the highest expression of proliferation, embryonic and oncogenic markers, and the lowest levels of differentiation markers.

We next investigated if GRN activity in the 369 HCC patients of the TCGA cohort correlated with prognosis. Fig. 1e shows that the survival probability as a function of time was lower in the 100 HCC patients with high GRN activity than in the 269 HCC patients with low GRN activity. When performing the same analysis with individual components of the GRN, we found that expression of *Let-7b, MYC* and *TP53* did not correlate with survival, while expression of *CTNNB1, SOX9, SMARCA4* and *LIN28B* were inversely correlated with survival (Supplementary Fig. 3). In this analysis of individual components, the cohort with high expression corresponded to the group of 100 patients with highest expression of the component, and the low expression cohort corresponded to the 269 other samples of the cohort. Together, the results indicate that the activity of the GRN as whole is a good marker of prognosis in HCC patients.

In conclusion, we have identified a GRN of functionally interacting partners which are misexpressed in HCC. The activity of the GRN, characterized by consistent and concomitant misexpression of its components and targets, correlates with proliferation, dedifferentiation and prognosis.

### The gene regulatory network promotes a proliferative HCC phenotype but is not correlated with telomerase maintenance

HCC’s are broadly divided in proliferative and nonproliferative classes ^1, 27^. Dedifferention and high proliferation characterize the proliferative class and correlate with the GRN activity (Fig. 1d), suggesting that the GRN is associated with or promotes a proliferative phenotype of HCC’s. This is further supported by our observation that GRN activity, defined as above, correlates with increased MET signaling, a pathway whose activation characterizes a subset of the proliferative HCC class ^28^. Indeed, HCC’s with low or high GRN activity and non tumor control samples from the TCGA cohort clustered separately in a PCA analysis when considering the expression of 110 HGF/MET target genes (Fig. 2a and Supplementary Fig. 1d). Moreover, to capture the tumors with highest HGF/MET target gene expression we selected 150 samples with the highest Dimension 1 and out of these we selected the 100 samples with highest Dimension 2 (Supplementary Fig. 1e). Those 100 tumor samples displayed increased expression of *CTNNB1*, *SOX9*, *SMARCA4*, *MYC* and *TP53* as compared to the other 269 tumors (Fig. 2b and Supplementary Fig. 1e).

**Figure 2.**
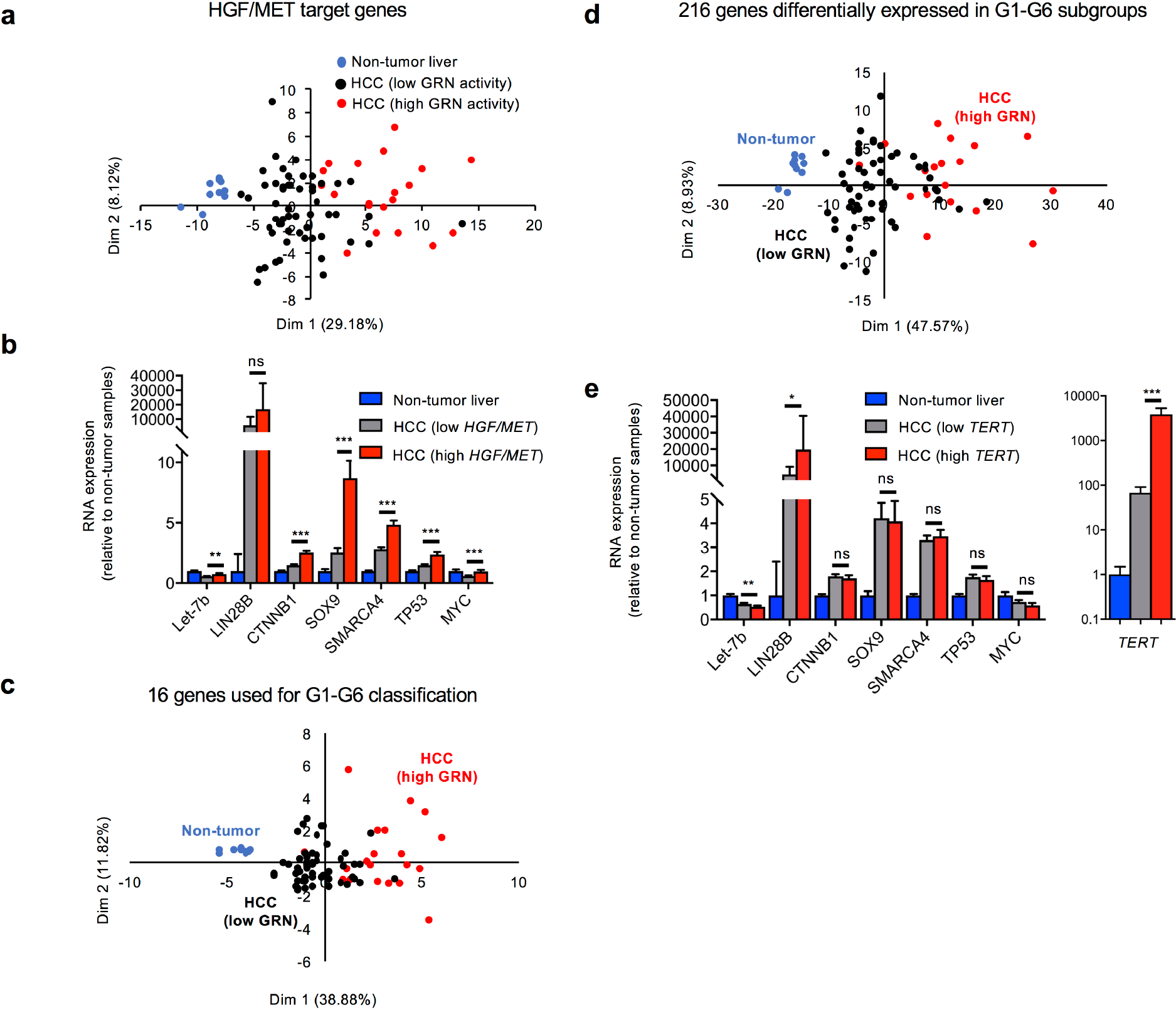
GRN activity correlates with HGF/MET pathway activation but not with telomerase expression. (**a**) Non-tumor samples (blue; n= 50) and HCC’s (n=369) cluster separately in a PCA analysis based on the expression of 110 HGF/MET target genes (see Table 1 in Ref. ^28^). Within the HCC samples, those with high GRN activity (red; GRN activity defined as in Fig. 1d) were separate from those with low GRN activity (black). (**b**) Expression of GRN components in non tumor samples (blue, n=50) and in HCC with low (grey; n=269) or high HGF/MET target expression (red; n=100). 100 tumor samples with high HGF/MET signaling were selected by PCA analysis based on the expression levels 110 of HGF/MET target genes (see panel a and Supplementary Fig. 1e). (**c, d**) Clustering of normal (blue) *versus* HCC with low (black) and high GRN activity (red) in a PCA analysis based on (**c**) the expression of 16 predictor genes and (**d**) of 216 differentially expressed genes of the 6-group HCC classification ^29^. Each dot in panels a, c and d represents the mean of 5 samples grouped in alphabetic order of the sample ID. (**e**) Expression levels of GRN components in non tumor (blue, n=50) and tumor conditions with low (grey, n=69) or high *TERT* expression (red, n=100). (**b, e**) Data are means +/− SEM. *, p<0.05; **, p<0.01 and ***p<0.001.

Earlier analyses correlated the transcriptome, genotype and phenotype of HCC patients and identified 6 patient subgroups numbered G1 to G6 ^29^. A predictor formula considered the expression of 16 genes to classify patients within one of the 6 subgroups, and, numerous genes were differentially expressed in the 6 subgroups ^29^. PCA analysis of the expression of a random selection of 216 differentially expressed genes or of the 16 predictor genes in non tumor tissue and HCC tumor of patients from the TCGA cohort revealed that tumors with high GRN activity cluster the farthest from the non tumor tissue (Fig. 2c-d and Supplementary Fig. 1f-g). Among the 216 genes differentially expressed between the G1 to G6 subgroups, the 50 genes that contribute the most to the Dimension 1 in the PCA plot are all upregulated in the G1, G2 and G3 subgroups and are mainly involved in cell proliferation (listed in Supplementary Table 2), fitting with our observations that GRN activity may contribute to confer a transcriptomic profile typical of the proliferation class of HCC. The G1, G2 and G3 subgroups are associated with poor differentiation, severe prognosis, and overexpression of genes regulating cell proliferation and DNA metabolism ^29^. Together, the data support that the GRN is associated with and may promote a proliferative phenotype.

*TERT* promoter mutation are found in 60% of HCC’s ^30^. However, out of the 369 HCC samples of the TCGA cohort, the 100 samples with highest TERT expression did not consistently misexpress GRN components as compared to the 269 HCC samples with lower *TERT* expression levels (Fig. 2e). This suggests that GRN activity is disconnected from *TERT* overexpression-induced telomere maintenance.

### Design of a mathematical model of the gene regulatory network

We next developed a tool to determine how variation of an individual GRN component affects the expression of all the others. To this end, we built a quantitative mathematical model, using a set of kinetic equations describing the expression of each network component as a function of time. The model includes MYC, TP53, SMARCA4, SOX9, LIN28B mRNA and protein, as well as active and inactive forms of CTNNB1 protein, which correspond to stabilized unphosphorylated CTNNB1 and phosphorylated CTNNB1 incorporated in a degradation complex, respectively. It also considers free (unbound) *Let-7b* miRNA, as well as complexes between *Let-7b* and mRNAs coding for SOX9, LIN28B and MYC. The variables, parameters and kinetic equations of the model are listed in Supplementary Tables 3-5.

We quantitatively calibrated the model using the mRNA levels of all GRN components available in TCGA (non tumor tissue and HCC tumors) and in a transcriptomic dataset from 34 human HCC cell lines ^31^ (Fig. 3a-c). Distinct sets of parameter values and initial conditions were determined for normal tissues on one hand, and for HCC tumors and cell lines on the other hand (Supplementary Tables 4 and 7).

**Figure 3.**
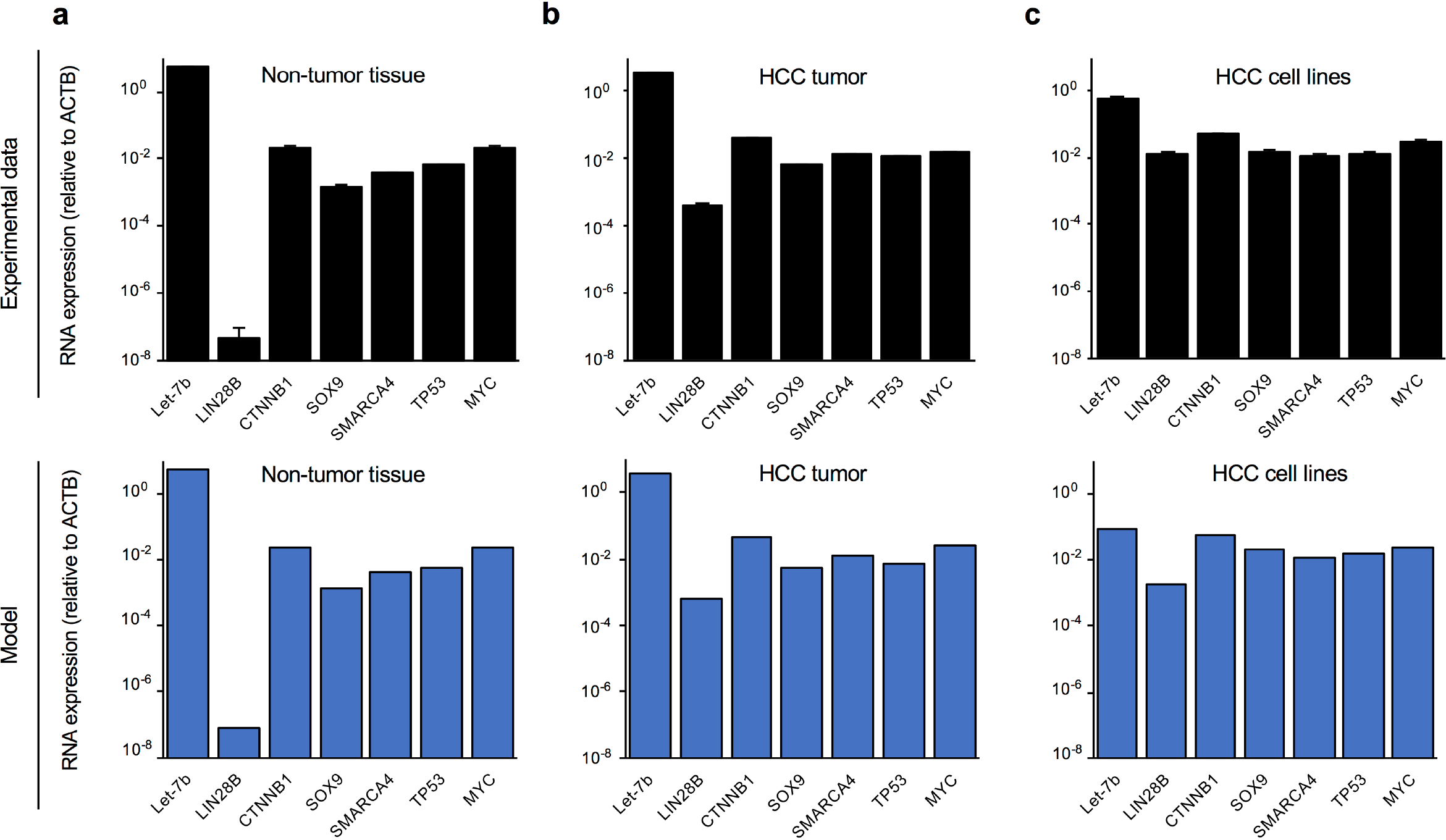
Calibration of the mathematical model on RNA expression levels in patient samples and human HCC cell lines. RNA levels of the GRN components are shown (**a**) in non tumor samples (n=50), (**b**) in HCC samples (n=369), and (**c**) in HCC cell lines (n=34). Data are means +/− SEM; patient data are from TCGA. The lower panels in a-c illustrate the expression of the GRN components in the three conditions as calculated using the mathematical model with the parameter and initial condition sets defined in Supplementary Tables 4 and 7.

Next, to quantitatively calibrate the model with protein expression values we transiently overexpressed GRN components in cultured HCC cell lines. These experiments validated in HCC the cross-regulating pairs of the GRN which had originally been identified in non-hepatic tumors. Overexpression of MYC in HepG2 or Huh7 cells stimulated TP53 and LIN28B expression, thereby validating in HCC the MYC→TP53 and MYC→LIN28B interactions (Figs. 4a, Supplementary Fig. 4a). Similarly, overexpression of constitutively active CTNNB1^S33Y^ and of SMARCA4 in HepG2 or Huh7 cells validated the CTNNB1→LIN28B and CTNNB1→MYC and SMARCA4→MYC interactions in HCC (Fig. 4b, Supplementary Fig. 4b-c). Finally, transient transfection of *Let-7b-5p* mimic RNA significantly repressed LIN28B, MYC and SOX9, confirming the regulatory links in the GRN (Fig. 4c).

**Figure 4.**
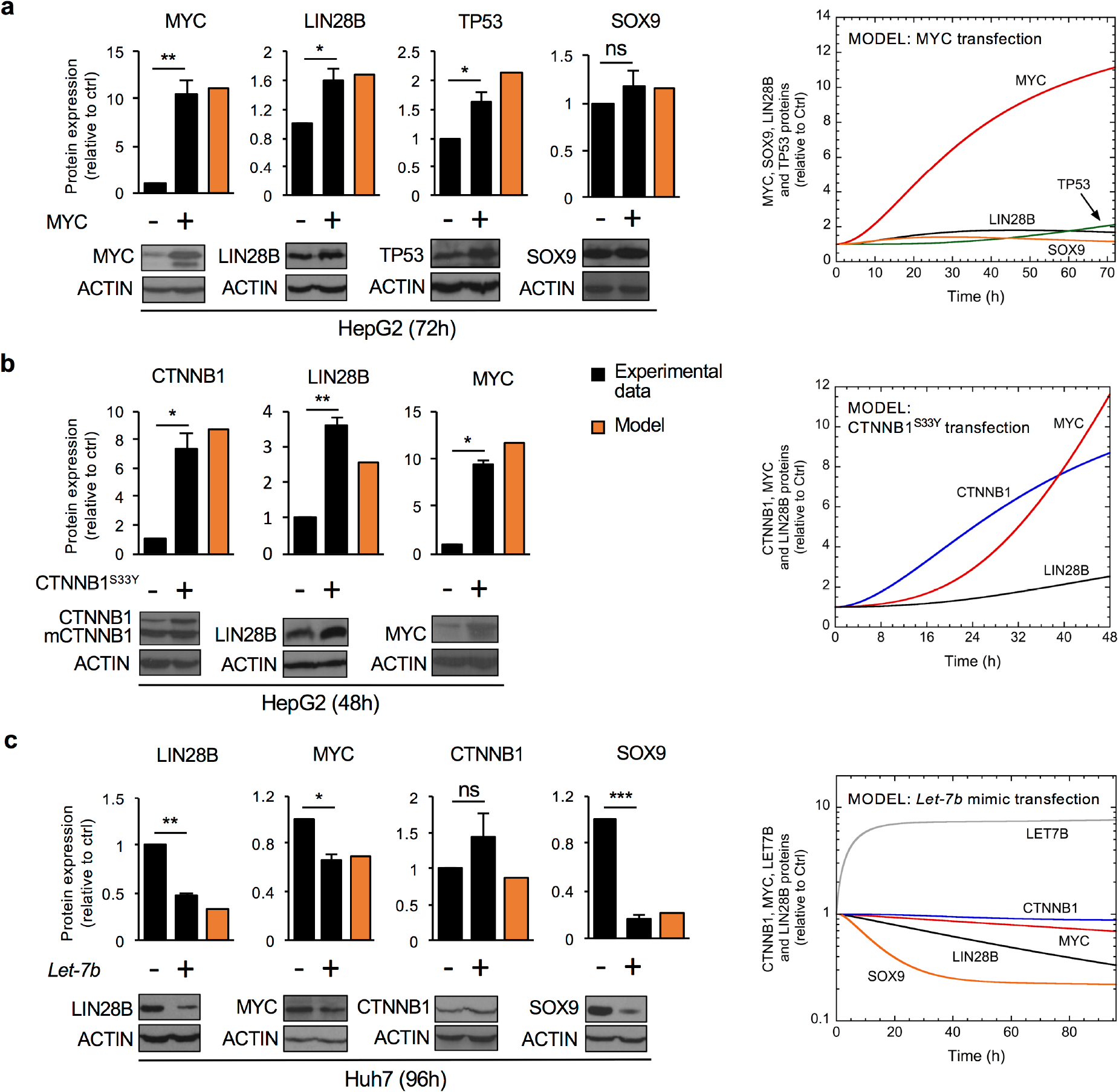
Calibration of the mathematical model on protein expression in human HCC cell lines. (**a**) Expression of MYC, LIN28B, TP53 and SOX9 protein following MYC overexpression in HepG2 cells. (**b**) Expression of CTNNB1, LIN28B and MYC protein following CTNNB1^S33Y^ overexpression in HepG2 cells. mCTNNB1 corresponds to endogenous mutant CTNNB1 resulting from inframe deletion of exon 3. (**c**) Expression of LIN28B, MYC, CTNNB1 and SOX9 in the presence of *Let-7b-5p* mimic RNA in Huh7 cells. Data in bar graphs are means +/− SEM, n ≥ 3. Orange bars correspond to protein levels calculated in the mathematical model with (**a**) overexpression of MYC (*V*_SMYC_ increases from 0.002 to 0.05 for 0 < t < 72h), (**b**) overexpression of CTNNB1^S33Y^ (for 0 < t < 48h *V*_S1CTNNB1_ increases from 0.0015 to 0.04 while *k*_ACTNNB_ varies from 0.05 to 10 and kICTNNB from 10 to 0.1) and (**c**) overexpression of *Let-7b* (for 0 < t < 96h, V_S1LET7_ increases from 0.01 to 0.1). The right panels in a-c show the simulated temporal evolution, with the appropriate time scale, of the GRN proteins following MYC, CTNNB1^S33Y^, or *Let-7b* mimic transfection. See Supplementary Information for details and Supplementary Tables 4 and 7 for parameter values and initial conditions for the variables used in the model.

The duration of the transient transfections in Fig. 4 (48h, 72h, 96h) reflected the minimal time-lapse required to monitor a significant change in TP53, LIN28B, MYC or SOX9 following overexpression of their respective stimulator or repressor. Therefore, both the level of protein induction and the timing required to observe a significant change in protein level were used to further calibrate the mathematical model of the GRN, while maintaining the quantitative calibration of all mRNA expression levels. We then simulated the transient transfection conditions by increasing the transcription rate constants of MYC, CTNNB1 or *Let-7b,* starting at t = 0h (Fig. 4a-c, right panels). The results showed that simulating a 10-fold increase in MYC -which mimics the observed 10-fold increase in transfected MYC protein- predicted a ~2-fold increase in LIN28B and TP53 after 72h; this predicted increase in LIN28B and TP53 matched closely the measured values (compare black and orange bars in Fig. 4a). Similarly, simulating an 8-fold increase in CTNNB1^S33Y^ predicted a ~3- and 11-fold induction of LIN28B and MYC after 48h; these inductions again matched closely the measured 4- and 9-fold induction of LIN28B and MYC, following an 8-fold increase in transfected CTNNB1^S33Y^ (Fig. 4b). Moreover, following a simulated 8-fold induction of *Let-7b* the model reproduced quantitatively the experimentally measured impact of *Let-7b* induction on LIN28B, MYC, and SOX9 (Fig. 4c). Finally, Let-7b induction in HuH7 cells did not affect *CTNNB1* expression, which fitted with the model simulation (Fig. 4c). We concluded that the calibrated mathematical model faithfully recapitulates the expression of GRN components.

### Validation of the mathematical model

To validate the model, we first challenged it by predicting the impact of a stabilizing CTNNB1 mutation on LIN28B and Let-7b expression and compared the prediction with values from the TCGA database. To this end we first partitioned the HCC samples of the TCGA database in two cohorts characterized by the presence or absence of stabilizing CTNNB1 mutation, the most frequent CTNNB1 mutations in the cohort being missense mutations in exon 3. This revealed that CTNNB1 mRNA levels were higher in the presence of a mutation (Fig. 5a, left). Therefore, to simulate CTNNB1 stabilization in the model we increased the *CTNNB1* mRNA synthesis rate (Supplementary information), increased the activation rate of the inactive (complexed) CTNNB1 form, and decreased the inactivation rate of the active (stabilized) form. Under these conditions, when *CTNNB1* mRNA values from the mutated and non-mutated samples were introduced in the model (Fig. 5a, orange bars), it faithfully predicted the reduction in *Let-7b* observed in non-mutated HCC samples as well as the slightly larger decrease of *Let-7b* seen in HCC samples with *CTNNB1* mutation. Also, the model predicted the increased *LIN28B* expression in HCC which was similar whether *CTNNB1* was mutated or not (compare orange and black bars in Fig. 5a).

**Figure 5.**
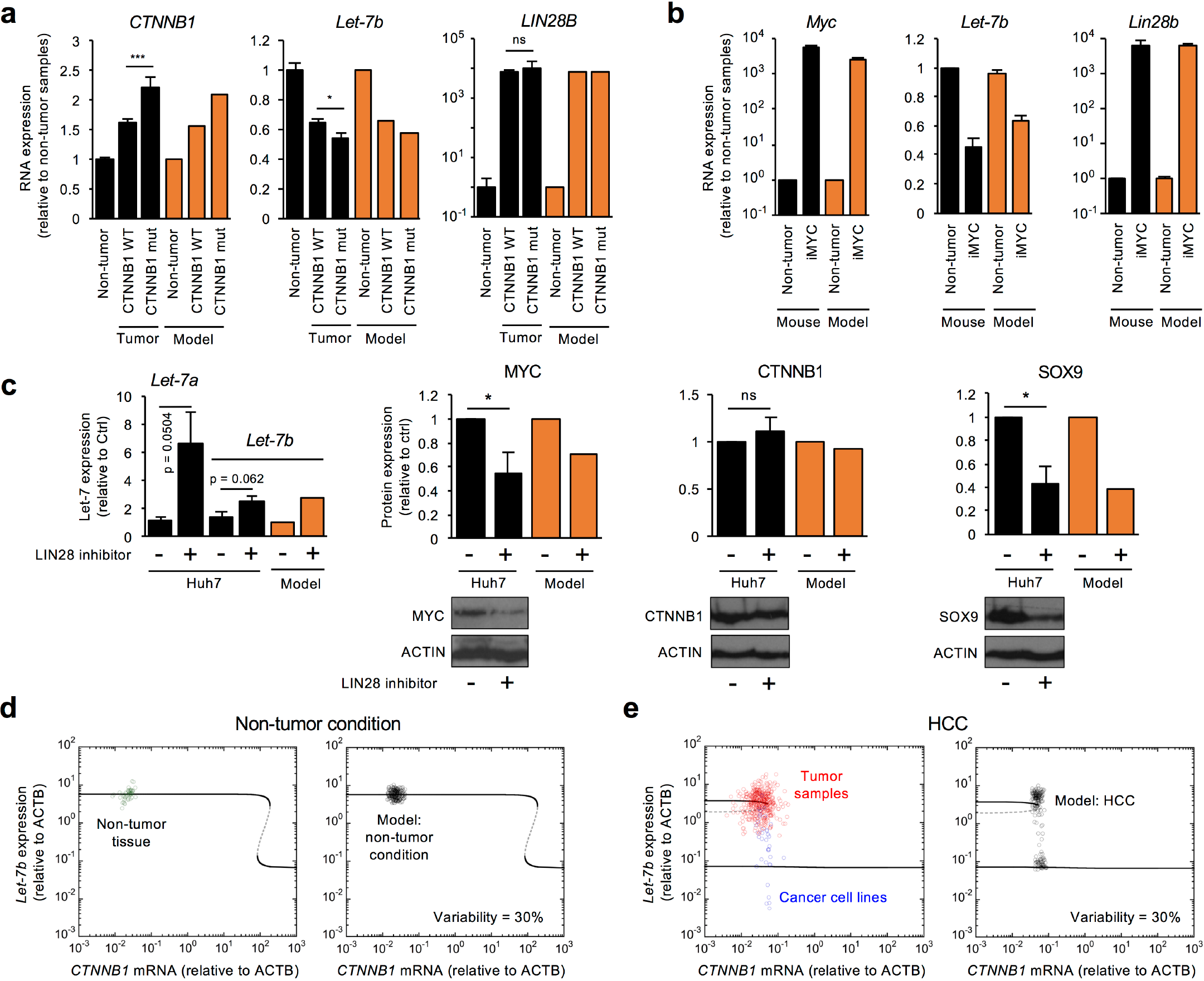
Validation of the model and GRN dynamics in HCC. (**a**) Expression of stabilized CTNNB1 mutant (n=98) is more increased than wild-type CTNNB1 (n=270) in HCC as compared to normal tissue (n=50). The predicted levels of *Let-7b* and *LIN28* following a simulation of increased expression and stabilization of CTNNB1 fit with the experimental observations. To simulate the stabilization of CTNNB1, the activation rate constant of CTNNB1 *k*_ACTNNB_ was raised from 1 (wild-type) to 10 (mutant), and the inactivation rate constant *k*_ICTNNB_ was decreased from 1 (wild-type) to 0.1 (mutant); other parameter values and initial conditions are in Supplementary Tables 4 and 7. (**b**) Measured and simulated impact of MYC induction on *LIN28B* mRNA and *Let-7b* expression levels validates the model. MYC induction was modeled by increasing V_SMYC_ from 0.002 to 6. Simulations were performed on a heterogeneous cell population of 200 cells with 30% of uniform random variations around the basal value of each parameter. (**c**) Simulating the absence (*K*_ILET7_ = 21) or presence (*K*_ILET7_ = 82) of LIN28 inhibition for 6 days (144h) predicts expression levels of *Let-7b,* and MYC, CTNNB1 and SOX9 which are close to the levels measured following treatment of Huh7 cells with 120 μM of LIN28 inhibitor *N*-Methyl-*N*-[3-(3-methyl-1,2,4-triazolo[4,3-*b*]pyridazin-6-yl)phenyl]acetamide. (**b, c**) Results are means +/− SEM; n ≥ 3 *, p<0.05. (**d**) Modelling *Let-7b* levels as a function of *CTNNB1* mRNA in normal conditions (left). Values in non tumor tissue (n=50; green dots) are superimposed on the model curve. Solid curves, stable steady states; dashed curves, unstable steady states. Modelling 30% random variations around the basal value of each parameter in 200 cells resulted in a distribution of *Let-7b/CTNNB1* values in the upper stable state (right) which recapitulated the heterogeneity measured in normal samples. (**e**) Modelling *Let-7b/CTNNB1* mRNA values in HCC conditions (left). *Let-7b/CTNNB1* mRNA values in 369 HCC tumors (red dots) or 34 cell lines (blue dots) are superimposed on the model curve: HCC patient tumors and cell lines are in distinct states. Random variation of parameters like in (d) recapitulated the heterogeneity measured in HCC patients and cell lines (right).

A second validation was obtained by considering previous results from a mouse model of MYC-induced liver cancer ^15^. We simulated a 4000-fold increase in *MYC* mRNA, which corresponds to the experimental induction of *Myc* in mice. Our simulation predicted a MYC-induced 7000-fold overexpression of *Lin28b* mRNA and a 0.4-fold reduction in *Let-7b.* This prediction fitted well with the experimentally-measured induction of *Lin28b* and reduction of *Let-7b* (Fig. 5b).

Third, as a proof of concept that the mathematical model can predict the impact of a pharmacological inhibitor of a GRN component, we simulated the inhibition of LIN28B in HCC: simulating 60 % inhibition of LIN28B protein predicted an increase in *Let-7b* and a decrease in SOX9 and MYC protein; CTNNB1 remained unaffected (Fig. 5c). We then evaluated the results of the simulation by growing Huh7 cells in culture in the presence of the LIN28 inhibitor (*N*-Methyl-*N*-[3-(3-methyl-1,2,4-triazolo[4,3-*b*]pyridazin-6-yl)phenyl]acetamide) ^32^. The inhibitor reduced cell proliferation (not shown), and, as expected, induced the expression of *Let-7b* and *Let-7a.* Importantly, the changes of SOX9, MYC protein and CTNNB1 protein levels measured by western blot matched well with the simulations (Fig. 5c), thereby validating the mathematical model as a tool to predict the impact of a pharmacological inhibitor on the GRN.

### GRN dynamics are characterized by a bistable switch

To characterize the dynamical properties of the GRN and determine whether they would identify distinct HCC cell states, we modelled the steady-state levels of *Let-7b,* selected as a representative variable, as a function of *CTNNB1* mRNA. In normal conditions, high *Let-7b* was associated with low *CTNNB1* mRNA, and *vice versa* (Fig. 5d, left). The system exhibits a reversible bistable switch from high to low levels of Let-7b, at supraphysiological levels of *CTNNB1* mRNA. Green dots in Fig. 5d (left) represent measured *Let-7b/CTNNB1* mRNA values in normal samples from the TGCA cohort. However, simulating *Let-7b* as a function of *CTNNB1* mRNA in HCC conditions revealed an irreversible bistable switch occuring at low *CTNNB1* mRNA levels (Fig. 5e, left). Indeed, rising *CTNNB1* mRNA from low to supraphysiological levels in HCC would induce a switch from high to low *Let-7* expression, but reverting from supraphysiological *CTNNB1* to low *CTNNB1* levels would not allow to restore high *Let-7* levels. Interestingly, individual *Let-7b/CTNNB1* values in HCC tumors clustered around the upper stable steady-state of the bistable switch (Fig. 5e, left, red dots). In contrast, blue dots in Fig. 5e represent *Let-7b/CTNNB1* values measured in 34 human HCC cell lines, and most are in the bottom stable steady state of the bistable switch, suggesting that culture conditions of cell lines promote switching of the GRN to a state distinct from that of the mean cell state in tumoral tissue of patients.

The distribution of measured *Let-7b/CTNNB1* values in Fig. 5d-e reflects intersample heterogeneity. When modelling a normal heterogeneous cell population (200 cells) with 30% of random variations around the basal value of each parameter, the *Let-7b/CTNNB1* values remained clustered in the upper stable steady state (Fig. 5d, right, black dots). Modelling a HCC cell population with the same % of parameter variation generated a distribution of *Let-7b/CTNNB1* values similar to that seen in HCC patients and cell lines (Fig. 5e, right), indicating that the model accounts for the sample’s heterogeneity.

We next analyzed the robustness of the bistable switch dynamics in normal and HCC conditions towards variation of parameter values. We plotted the steady-state levels of *Let-7b* as a function of *CTNNB1* mRNA levels in the presence of a two-fold increase or decrease of each parameter value (Supplementary Figs. 5 and 6). In normal condition, *i.e.* high *Let-7b* and low *CTNNB1* mRNA, the model is very robust towards 2-fold parameter variation since high *Let-7b* levels are maintained regardless of parameter variations (Supplementary Fig. 5). In contrast, in HCC conditions the irreversible bistable switch dynamics can be lost following parameter variations, and only the bottom stable steady state of the bistable switch is maintained (Supplementary Fig. 6). Thus, the tumor state is more sensitive towards parameter variations than the normal state. This is expected to be reflected in patient samples by increased heterogeneity in expression of the GRN components, as observed in Fig. 1b.

We concluded that the GRN dynamics rest on a robust reversible bistable switch that should not occur in normal condition, while it exhibits a sensitive irreversible bistable switch in HCC condition that could cause heterogeneity among tumor samples.

### Gene regulatory network activity in gastrointestinal cancers

Since the components of the GRN are expressed in several tissues we verified if it might be activated in cancers distinct from HCC. We first looked at RNA expression in TCGA cohorts of cholangiocarcinoma (CHOL), stomach and esophageal carcinoma (STES), colorectal adenocarcinoma (COADREAD), thyroid carcinoma (THCA), kidney renal clear cell carcinoma (KIRC), breast carcinoma (BRCA), bladder urothelial carcinoma (BLCA), and lung adenocarcinoma (LUAD) (Fig. 6a, Supplementary Fig. 7). Statistically significant and consistent induction of all tumor-promoting GRN components was detected in cholangiocarcinoma, stomach and esophageal carcinoma, and colorectal adenocarcinoma, *i.e.* in gastrointestinal cancers. *Let-7b* did not show the expected reduction in those tumors, yet depending on the tumor type, other *Let-7* family members are downregulated, like in cholangiocarcinoma where *Let-7c* is strongly repressed. The other cancer types did not show a consistent increase in GRN components, suggesting that the GRN does not display tumor-promoting activity in these tumors.

**Figure 6.**
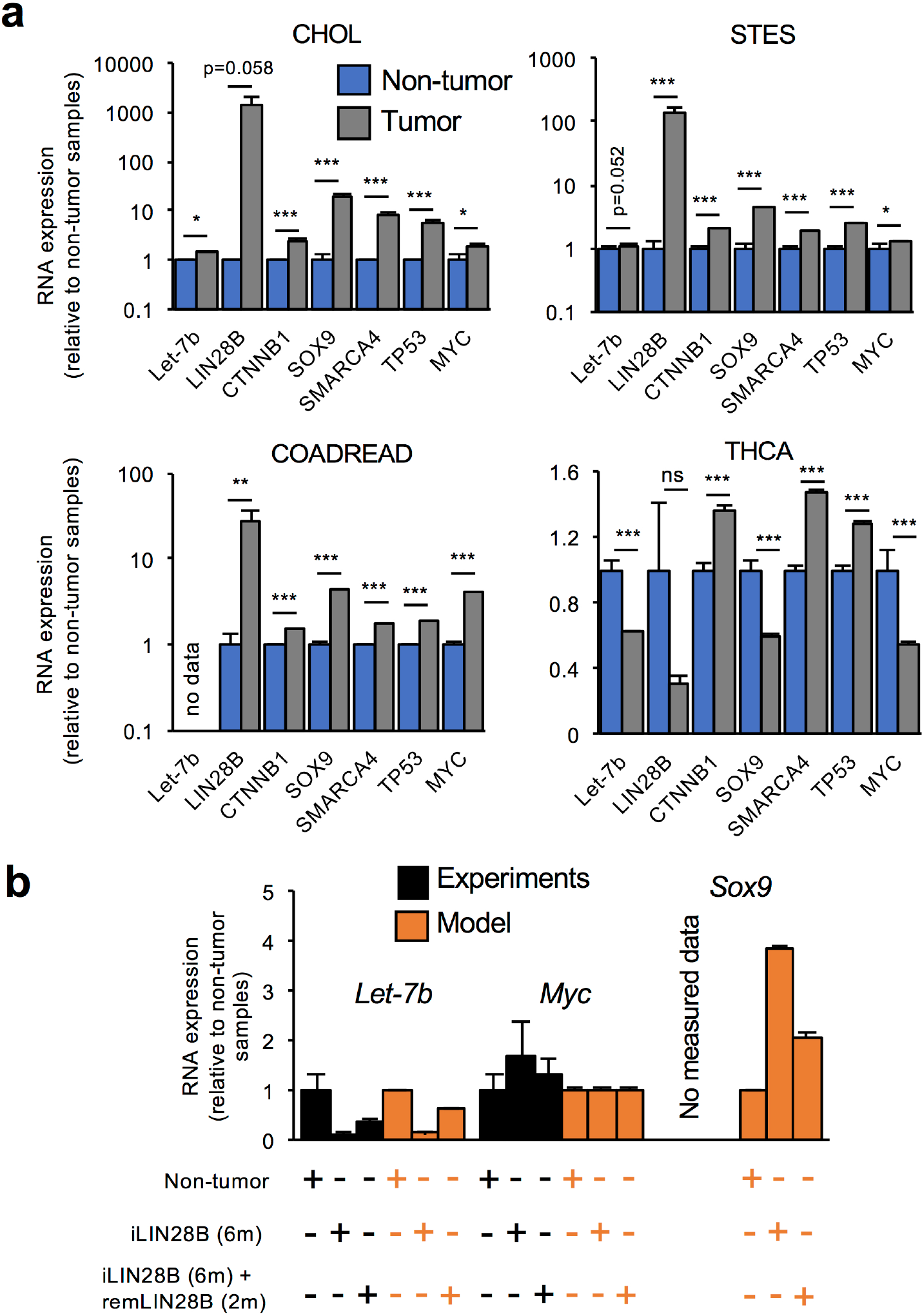
Cancer-type specificity of GRN activity and validation of the model in colorectal cancer. (**a**) Tumor-promoting components of the GRN are induced in cholangiocarcinoma (CHOL), stomach and esophageal carcinoma (STES), and colorectal adenocarcinoma (COADREAD). Thyroid carcinoma (THCA) had a distinct gene expression profile. Data (mean +/− SEM) are from TCGA; the number of normal and tumor samples is 9 and 36 (CHOL), 50 and 600 (STES), 51 and 624 (COADREAD), and 59 and 501 (THCA). *, p<0.05, **, p<0.01 and ***p<0.001. (**b**) RNA levels of *Let-7b, Myc* and *Sox9* in experiment (black) and in the mathematical model (orange) in non tumor condition, after 6 months (4320 h) of *Lin28b* induction (iLIN28B; *tumor* = 1 and *xLIN28* = 0.005; Supplementary Table 4), and after 6 months of *Lin28b* induction followed by 2 months of partial *Lin28b* removal (remLIN28B; *tumor* = 1 and *xLIN28* = 0.0015). The predicted expression levels are means +/− SEM of a heterogeneous cell population with 30% of uniform random variation around the basal value of each parameter. The modelled cell population consists of 200 cells with stabilizing *CTNNB1* mutation (*k*_ACTNNB_=10 and *k*_ICTNNB_=0.1) and 200 cells without *CTNNB1* mutation (μACTNNB=0.05 and AICTNNB=10). All parameter values used in each condition are in Supplementary Information.

Tu and coworkers developed a mouse model of colorectal cancer with similar gene expression pattern as in human cancer ^21^, which offered the opportunity to validate the mathematical model in this cancer type. The published mouse data showed that induction of *Lin28b* for 6 months triggered cell proliferation, downregulation of *Let-7* and upregulation of *Sox9,* the latter being illustrated by immunostaining ^21^. The subsequent removal of *Lin28b* for 2 months partially restored the expression of *Let-7* and proliferation markers. As in the experiments ^21^, the mathematical model predicts that a 6-month induction of *LIN28B* leads to downregulation of *Let-7b* and upregulation of *SOX9,* and that those expression levels partially revert to near normal values 2 months after *LIN28* removal (Fig. 6b), thereby validating the GRN model in colorectal cancer. The mathematical model also shows that the expression of *MYC* (Fig. 6b) and *CTNNB1* mRNAs (not shown) is not affected by *LIN28B* induction, which fits well with the observations ^21^.

## Discussion

The pattern of genetic and epigenetic alterations is heterogeneous in HCC. Therefore, to understand the mechanisms of disease progression and design therapeutic strategies, there is a need to identify the functionality and the dynamics of GRNs driving HCC in individual samples ^4, 29, 33^. Here we identified a GRN consisting of functionally-interacting components and which is activated in HCCs and gastrointestinal tumors. The GRN combines CTNNB1, SMARCA4, SOX9, LIN28B, Let-7b, TP53 and MYC, which are misexpressed and/or mutated in HCC. *MYC* expression is not strongly affected in the HCC cohort but it is higher in tumors with high GRN activity than in tumors with low GRN activity, and plays a role in mouse models of HCC ^15^.

Since the mathematical model resorts to two sets of parameters, one for normal condition and one for tumors and cell lines, it does not account for the dynamics of transition between normal and HCC states. Indeed, to properly calibrate the model on the mRNA expression levels, the transcription rates of *LIN28B, SOX9,* mutant *TP53, CTNNB1* and *SMARCA4* had to be increased in the HCC conditions (see *‘tumor’* parameter in Supplementary Tables 4 and 5). The need to increase these parameters indicates that the interactions between GRN components are not sufficient to account for the normal-to-HCC transition. The model parameters implicitly integrate the impact of external regulators of the GRN. Yet, the modelling strategy cannot integrate the full spectrum of regulations, and a number of regulators might not be known. Since modelling the transition from normal to HCC requires adaptation of the transcription rates of *LIN28B, SOX9,* mutant *TP53, CTNNB1* and *SMARCA4,* we suggest that the mechanisms controlling the expression of those genes warrant further investigation.

PCA analysis based on the expression of GRN components allowed us to define subclasses of patients. Molecular classifications of HCC’s identified proliferative *versus* non proliferative classes, and another approach correlated the transcriptome, genotype and phenotype of HCC patients to subdivide patients in 6 subgroups called G1 to G6 ^29^. Our observations suggest that GRN activity may contribute to confer a transcriptomic profile typical of the proliferative class of HCC. Consistently, the GRN is most activated in the G1, G2 and G3 subgroups which are associated with poor differentiation, severe prognosis, and overexpression of genes regulating cell proliferation ^29^. The targets of CTNNB1 are heterogeneous as it induces progenitor-type genes in tumor cells such as cyclin-D1 or VEGF-A, but also regulates expression of antioxidant, pro-survival and pro-hepatocyte differentiation genes in normal hepatocytes ^34^. The G5 and G6 HCC subgroups are associated with activation of CTNNB1 targets typical for mature hepatocytes, whereas the G1, G2, and G3 subgroups show predominant activation of progenitor-type targets. Interestingly, high GRN activation is associated with high expression of progenitor-type CTNNB1 targets (Supplementary Fig. 8).

The structure of the GRN comprises several positive feedback loops which are at the origin of reversible and irreversible bistable switches. Bistable switches have been identified in multiple molecular regulatory networks involved in differentiation, signaling or proliferation pathways ^35, 36, 37, 38^. However, our quantitative modelling approach identified the first irreversible bistable switch involved in a specific subset of HCC tumors. It sheds light on the nature of a dynamical mechanism which can be a source of heterogeneity among HCC samples. In addition, the identification of irreversible states in HCC provides evidence that targeting specific GRN components using drugs might be therapeutically ineffective, given that the cancer cell is in a locked state with regard to the function of the target. Also, our comparison of patient tumors and cultured HCC lines (Fig. 5e) indicates that cell lines may be in a state distinct from that of tumors with regard to the function of the GRN. This raises the need for caution when testing the efficacy of drugs in cultured cells. To facilitate the analysis of the GRN, we set up a user-friendly web-platform allowing to check GRN activity in HCC and in gastrointestinal tumors. This platforms also presents a graphical user interface that integrates the expression levels of the GRN components of new tumor samples, and which implements the mathematical model to test which component of the network is the best target to modulate the network dynamics (http://biomodelling.eu/apps.html). We anticipate that identification and modelling of a set of GRNs consisting each of functionally-interacting components will significantly contribute to provide a global picture of tumor-promoting gene function in HCC. Our study presents the concept of tools that help in the design of bespoke therapies for treating each patient’s specific cancer.

## Methods

### Description and calibration of the mathematical model

The mathematical model of the GRN is based on 20 kinetic equations describing the temporal evolution of the expression level of each network component (Supplementary Information). The variables, parameters, initial conditions and numerical values used to calibrate the model are mentioned in Supplementary Tables 3-7.

### Data normalization and statistical analysis

Data normalization methods are described in Supplementary Information. All measured data are means ± SEM. Significance was assessed by Student t-test.

### Mathematical modelling and PCA analysis

Mathematical model simulations were performed using XPPAUTO (http://www.math.pitt.edu/~bard/xpp/xpp.html) and Matlab. PCA analysis was performed with FactoMineR (R package) ^39^.

### RNASeq and miRNASeq data

RNASeq and miRNA Seq data of patient cohorts were from TCGA database http://firebrowse.org/). For each cohort, we converted the “scaled_estimate” in the “illuminahiseq_rnaseqv2_unc_edu_Level_3_RSEM_genes” file into TPM by multiplying by 10^6^.

### Plasmids and microRNAs

Plasmids pCDNA3.1, pcDNA3-MYC, pCl-neo β-Catenin (CTNNB1^S33Y^), pBABE-BRG1 were from Thermo Fisher Scientific (Waltham, MA, USA), Wafik El-Deiry (Addgene plasmid # 16011) ^40^, Bert Vogelstein (Addgene plasmid # 16519) ^41^ and Robert Kingston (Addgene plasmid # 1959) ^42^ respectively. The miR mimic control (MIMAT0000039: UCACAACCUCCUAGAAAGAGUAGA) and miR mimic hsa-let-7b: hsa-let-7b-5p (MIMAT0000063: UGAGGUAGUAGGUUGUGUGGUU), hsa-let-7b-3p (MIMAT0004482: CUAUACAACCUACUGCCUUCCC) were from Dharmacon (Lafayette, CO, USA).

### Cell culture and transfections

Human Huh7, HepG2 and Hep3B hepatocarcinoma cells were grown in DMEM (Lonza, Westburg, Leusden, Netherlands), 10% Foetal bovine serum (Merck, Darmstadt Germany), L-Glutamine (2 mM) (Thermo Fisher Scientific, Waltham, MA, USA), Penicillin-Streptomycin (50 U/mL and 50 μg/mL respectively) (Gibco™, Waltham, MA, USA) and Amphotericin B (Gibco™, Waltham, MA, USA) (2.5 μg/mL). Cells were grown in 60 mm dishes and transfected with 3 μg plasmid and 120 nM miR mimic, using jetPRIME^®^ (Polyplus-Transfection, Illkirch-Graffenstaden, France) for 48, 72 or 96h as recommended by the manufacturer, in at least three independent experiments. DNA was transfected 24h after plating the cells on 60 mm dishes. For LIN28 inhibition, Huh7 cells were grown in the presence of 120 μM of (*N*-Methyl-*N*-[3-(3-methyl-1,2,4-triazolo[4,3-*b*]pyridazin-6-yl)phenyl]acetamide) for 6 days; the medium with inhibitor was changed every day

### Protein extractions and western blotting

Cells were rinsed twice with PBS and total proteins were extracted using a cell lysis buffer (50 mM Tris-Cl pH 7.4, 150 mM NaCl, 1mM EDTA, 0.5% IGEPAL, 1mM DTT and protease inhibitors (Roche, Bâle, Switzerland). Cells were sonicated for 15s and centrifuged. Protein concentration was measured with a Bradford assay. Proteins (40 μg) were fractionated using 8-10% polyacrylamide gels, transferred to PVDF membranes (Merck, Darmstadt, Germany), and detected by enhanced chemiluminescence (ECL, Thermo Fisher Scientific, Waltham, MA, USA) using X-ray films (Thermo Fisher Scientific, Waltham, MA, USA) and Fusion Solo S equipment (Vilber Lourmat, Collegien, France). Proteins were quantified using Bio1D advanced software (Vilber Lourmat, Collegien, France). Antibodies and their dilutions were rabbit polyclonal anti-LIN28B antiserum (#4196S, 1:1000, Cell Signalling Technology), rabbit polyclonal anti-MYC antiserum (sc-764, 1:500, Santa Cruz Biotechnology), mouse monoclonal anti-CTNNB1 antibody (BD610154, 1:2000; BD Transduction Laboratories™), rabbit polyclonal anti-SOX9 antiserum (AB5535, 1:1000, Merck), mouse monoclonal anti-TP53 antibody (sc-126, 1:1000, Santa Cruz Biotechnology), rabbit polyclonal anti-BRG1 antiserum (sc-10768, 1:500, Santa Cruz Biotechnology) and goat polyclonal anti-ACTIN antiserum (sc-1615, 1:500, Santa Cruz Biotechnology).

### RNA extraction and analysis

Total RNA was isolated from Huh7 cultured cells, in the presence or absence of LIN28 inhibitor for 6 days, using Trizol (#1029602, Invitrogen, Life technologies). cDNA synthesis was performed with MMLV reverse transcriptase (#28025-13, Invitrogen, Life technologies) according to manufacturer’s protocol. MicroRNA expression (Let-7a and Let-7b) was quantified by RT-qPCR using Kapa SYBR Fast 2X Universal Master Mix (#KK4601, Sopachem, Ochten, Netherlands). Specific stem-loop primers were used for reverse transcription, and RT-qPCR was performed using a specific forward primer and a common universal reverse primer. *Let-7a* Fwd/Rev: ACACTCCAGCTGGGTGAGGTAGTAGGTTG/CTCAACTGGTGTCGTGGAGTCGGC AATT CAGTT GAGACTATACA; *Let-7b* Fwd/Rev: ACACTCCAGCTGGGTGAGGTAGTAGGTTGT/CT CAACT GGTGTCGTG GAGT CGGCAATT CAGTT GAGAACCACAC; *ACTB* Fwd/Rev: TCCTGAGCGCAAGTACTCTGT/CTGATCCACATCTGCTGGAAG. In all conditions, each ΔCt between the measured transcripts and the housekeeping genes was further normalized to their control conditions by using 2^-ΔΔCt^ method.

## Acknowledgements

The authors thank P. Jacquemin, C. Pierreux and the Lemaigre laboratory members for help and input. We also thank L. Nguyen and H. Zhu for sharing information, and Lieven Desmet for his help with statistical analyses. The work of FPL was supported by the Interuniversity Attraction Pole Programme (Belgian Science Policy, PVII-47), the D.G. Higher Education and Scientific Research of the French Community of Belgium (ARC 15/20-065), the F.R.S.-FNRS (Belgium: Grants T.007214 and J.0058.15), and the Belgian Foundation Against Cancer (grant 2014-125). J.Z-R’s group is supported by INSERM, the Ligue Nationale contre le Cancer (Equipe Labellisée), Labex OncoImmunology (investissement d’avenir), Coup d’Elan de la Fondation Bettencourt-Shueller, the SIRIC CARPEM and Fondation Mérieux.

## Author contributions

C.G. and F.P.L. conceived and designed the study. C.G. performed transcriptomic data analysis and mathematical modelling. M. D-L., K.K. and S. Cordi performed transfection experiments and quantified RNA and proteins. L.G. created the web-based platform under supervision of E.H. A.L. did the survival analysis and helped with transcriptomic data analysis. S. Caruso, G.C. and J.Z-R. provided transcriptomic data and human cell lines and contributed to the interpretation of the modelling results. J.T. and S.P.M. shared and analyzed data from transgenic mice. C.G., M. D-L. and F.P.L. wrote the manuscript with input from all authors.

